# Efficient use of harvest data: An integrated population model for exploited animal populations

**DOI:** 10.1101/776104

**Authors:** Marlène Gamelon, Éric Baubet, Aurélien Besnard, Jean-Michel Gaillard, Jean-Dominique Lebreton, Laura Touzot, Lara Veylit, Olivier Gimenez

## Abstract

1. Many populations are affected by hunting or fishing. Models designed to assess the sustainability of harvest management require accurate estimates of demographic parameters (e.g. survival, reproduction) hardly estimable with limited data collected on exploited populations. The joint analysis of different data sources with integrated population models (IPM) is an optimal framework to obtain reliable estimates for parameters usually difficult to estimate, while accounting for imperfect detection and observation error. The IPM built so far for exploited populations have integrated count-based surveys and catch-at-age data into ageclass structured population models. But the age of harvested individuals is difficult to assess and often not recorded, and population counts are often not performed on a regular basis, limiting their use for the monitoring of exploited populations.
2. Here, we propose an IPM that makes efficient use of data commonly collected in exploited marine and terrestrial populations of vertebrates. As individual measures of body mass at both capture and death are often collected in fish and terrestrial game species, our model integrates capture-mark-recapture-recovery data and data collected at death into a body mass-structured population model. It allows the observed number of individuals harvested to be compared with the expected number and provides accurate estimates of demographic parameters.
3. We illustrate the usefulness of this IPM using an emblematic game species distributed worldwide, the wild boar *Sus scrofa*, as a case study. For this species that has increased in distribution and abundance over the last decades, the model provides accurate and precise annual estimates of key demographic parameters (survival, reproduction, growth) and of population size while accounting for imperfect detection and observation error.
4. To avoid an overexploitation of declining populations or an under-exploitation of increasing populations, it is crucial to gain a good understanding of the dynamics of exploited populations. When managers or conservationists have limited demographic data, the IPM offers a powerful framework to assess population dynamics. Being highly flexible, the approach is broadly applicable to both terrestrial and marine exploited populations for which measures of body mass are commonly recorded and more generally, to all populations suffering from anthropogenic mortality causes.

## 1 INTRODUCTION

Many animal populations are affected by commercial, recreational or subsistence harvest (Lebreton 2005b; Peres 2010; Ripple *et al.* 2016), i.e. by the removal of individuals through hunting or fishing. Managing such populations to keep the harvest at sustainable levels has long been a central purpose (Williams, Nichols & Conroy 2002). It is especially true in the current context of global change, as both experimental and observational evidence that harvest may act in synergy with other negative influences such as habitat destruction or disease outcome is accumulating (Camilo *et al.* 2007; Koons *et al.* 2015; Chen *et al.* 2015). Likewise, in the so-called harvest-interaction hypothesis, harvest might interact with population-level effects of climate change in both marine and terrestrial ecosystems. This interplay between harvest and climate effects may amplify environmentally induced fluctuations in population size and increases extinction risk, or, alternatively, dampen fluctuations and increase population growth rates (Gamelon, Sandercock & Sæther 2019).

Models designed to assess the sustainability of harvest management typically require accurate estimates of demographic parameters (e.g. survival, reproduction) and population size, which cannot be reliably estimated with limited data. Hence, in particular when demographic information is limited, the challenge is to make efficient use of available data to gain a good understanding of the dynamics of exploited populations and be able, in turn, to provide appropriate management recommendations. When several data types are available, even if each data type by itself provides limited information on demographic parameters, a combined analysis within an integrated population model (IPM) approach offers several advantages (see Schaub & Abadi 2011; Zipkin & Saunders 2018 for reviews). First, a combined analysis of different data sources always increases the precision of demographic estimates (for a proof, see Barker & Kavalieris 2001, and for an example, see table 1 in Péron *et al.* 2010 who compare parameter estimates and standard errors from a capture-mark-recapture only and an integrated-modeling approach). Second, imperfect detection and observation error inherently associated with data sampled in the field (e.g. population counts) are accounted for. Third, the use of IPM allows some parameters that are difficult or impossible to estimate based on separate analyses to be satisfactorily estimated (for an example, see Péron *et al.* 2010).

From the eighties, the simultaneous analysis of different data sources with IPM has received growing interest in fisheries (see Maunder & Punt 2013 for a review). Later, IPM have been proposed as powerful tools to assess the dynamics of terrestrial vertebrate populations (e.g. Besbeas *et al.* 2002 for a study on northern lapwing *Vanellus venellus* and grey heron *Ardea cinerea*) and more recently, they have been applied to exploited populations in terrestrial ecosystems (Gauthier *et al.* 2007; Arnold *et al.* 2018). Strikingly, IPM built on exploited populations usually integrate population surveys (of alive individuals), catch-at-age data and capture-mark-recapture-recovery (CMRR) data into age-structured population models (Methot Jr & Wetzel 2013; Arnold *et al.* 2018; Scheuerell *et al.* 2019). This state-of-the-art limited up-to-now the applicability of IPM for two reasons: i) the age of harvested individuals is often not available because its determination is challenging and generally involves expensive and time-demanding analyses. Instead, body mass at capture and/or death are commonly recorded; ii) information on numbers is rarely based on surveys of alive individuals. Instead, the number of individuals removed through exploitation is commonly recorded in many exploited species.

Here, we develop a widely applicable IPM relying on data commonly collected in exploited populations. We illustrate the usefulness of this approach based on the study of a population of an emblematic game species distributed worldwide, the wild boar *Sus scrofa*. This species has tremendously increased in abundance and distribution over the last decades (Massei *et al.* 2015), leading to important damage to crops and high risk of disease transmission in Europe (see e.g. Schulz *et al.* 2019). Hunting is commonly assumed to help controlling wild boar expansion. Our model differs from previous IPMs applied to vertebrate populations in two respects: i) typical IPMs integrate data into an age-class structured population model. Instead, we build an IPM that integrates CMRR data and individual measures recorded at death into a body mass-structured population model; ii) both alive and dead individuals are considered in the model allowing us to compare the observed number of individuals shot by hunters in each body mass class to the numbers expected from the IPM. For each mass class, the model allows us to get accurate and precise annual estimates of demographic parameters (i.e. survival, reproduction) and of the number of alive individuals, while accounting for imperfect detection and observation error. Although we targeted wild boar as a case study in this work, the approach we propose can be reliably used for assessing population dynamics of any exploited size-structured population of vertebrates.

## 2 MATERIALS AND METHODS

### 2.1 Demographic data collection

We studied a wild boar population located in the 11,000 ha forest of Châteauvillain-Arc-en-Barrois in North-Eastern France (48°02’N, 4°55’E). Between 1991 and 2016, thanks to an intensive capture-mark-recapture program, 1,152 females were captured from March to September using traps, marked and released in their environment. For each capture event, we recorded the date of capture and body mass. Between October and February, wild boars were harvested. All females shot by hunters (previously marked or not) were weighed and their date of death was accurately recorded. No information was available for individuals that died from natural causes. As wild boar rut generally begins in mid-December, females are often pregnant when shot during the hunting season. For pregnant females, the number of fetuses present in the uteri was recorded.

Three types of demographic data were thus available: CMRR data, hunting data and reproduction data.

i. CMRR data provided individual histories of 1,152 marked females together with the body mass at each capture (alive) and when shot (dead).
ii. Hunting data were the number of females shot by hunters in year *t, y*_*t*_ together with their body mass at death (n=7,350 over the study period).
iii. Reproduction data were the number of pregnant females shot in year *t, R*_*t*_ (n = 811 over the study period) together with their body mass as well as the number of fetuses at *t, J*_*t*_ (n = 4,344 over the study period).

The Integrated Population Model (see Fig. 1 for a schematic representation of the IPM we built) was made of two types of components (Schaub & Abadi 2011; Zipkin & Saunders 2018): i) a population model iterating a population vector, which links demographic rates (e.g. survival, reproduction) to the population vector; ii) probabilistic models and their likelihood functions for each of the three data sets (CMRR data, hunting data, reproductive data) separately. In turn, we obtained a joint likelihood combining these components.

**Figure 1.**
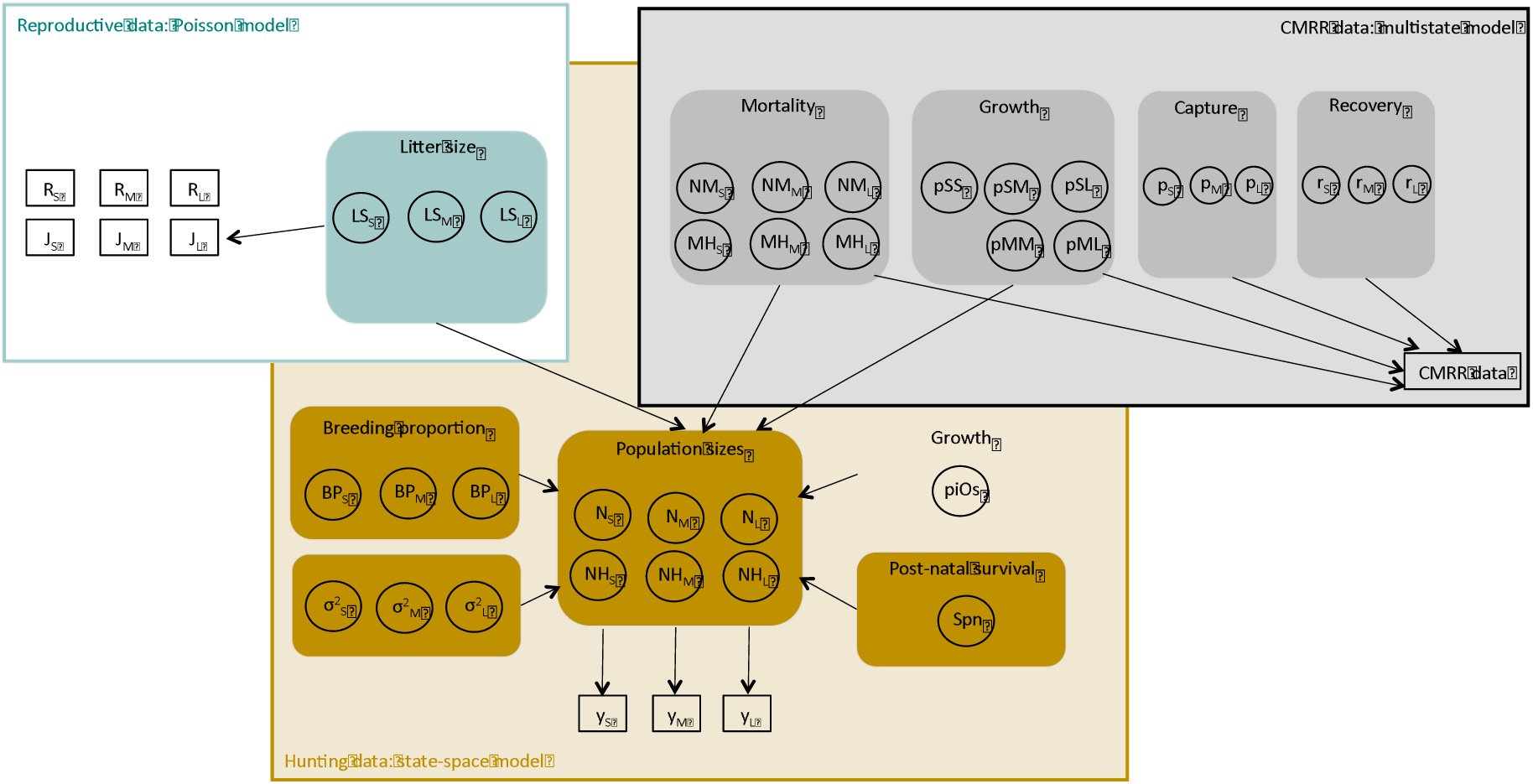
Directed acyclic graph of the IPM. Squares represent the data, circles represent the parameters to be estimated. Arrows represent dependencies. Three types of data are collected: CMRR data, hunting data (*y*_*j*_) and reproductive data (number of pregnant females (*R*) and number of fetuses (*J*) in each body mass class *j*). Estimated parameters are the body mass-specific recapture probability *p*_*j*_, recovery probability *r*_*j*_, natural mortality *NM*_*j*_, hunting mortality *MH*_*j*_, litter size *LS*_*j*_, numbers of females alive *N*_*i*_, numbers of females shot by hunters *NH*_*j*_, breeding proportions *BP*_*j*_, post-natal survival *Spn* (from birth to weaning), observation error 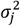 and transitions between body mass classes (growth) with *piOs* the probability for newborn remaining in the small body mass class, *pSS* the probability of small females remaining in this class during the year, *pSM* the probability of small females entering the medium class, *pSL* the probability of small females entering the large class, *pMM* the probability of medium females remaining in this class during the year and *pML* the probability of medium females entering the large class.

### 2.2 Population model

The population vector was considered each year at the end of the hunting season and before reproduction (pre-breeding census). From life history of wild boar and available data (data on body mass, not on age and data on numbers of individuals killed, not on numbers of alive individuals), the population vector considered three classes of body mass (<30 kg (small), between 30 and 50 kg (medium) and >50 kg (large) (see Gamelon *et al.* 2012)) and two states (alive and shot by hunters) for each body mass-class. The population vector had thus 6 components: 3 body mass-classes × 2 states.

The stage-structured matrix population model allows stage-specific population sizes at year *t+1* to be estimated from stage-specific population sizes at year *t* and demographic rates (Caswell 2001). We built a female body mass-structured matrix model by considering the three classes of body mass previously defined and the two states: alive and shot by hunters. The output of this body mass-structured matrix model structured according to body mass can be directly compared with the observed distribution of hunted individuals among body mass-classes. Females can reproduce in all the three classes. Females may remain in the same body mass-class from one year to the next, with a probability *pSS* for small females and *pMM* for medium females. All large females remained in the large body mass class. Alternatively, small females can move to the medium body mass class with a probability *pSM* or move directly to the largest class with a probability *pSL* (such as *pSL=1-pSS-pSM*). Medium females can move to the next class with a probability *pML* (such as *1-pMM*). There was no backward transition towards a lighter body mass class. As wild boar females are sedentary (Truvé & Lemel 2003; Keuling *et al.* 2010), we assumed no immigration and emigration.

We defined *N*_*S,t*_ as the number of small females, *N*_*M,t*_ the number of medium females and *N*_*L,t*_ the number of large females alive in year *t*. To account for demographic stochasticity in survival processes, we used binomial processes to describe the number of females in each body mass class.

i.The number of small females alive in year *t N*_*S,t*_ is the number of small females in year *t-1* that remained small plus the number of small females produced by mature females in year *t-1 (NewBorn*_*t-1*_) that survived, such as:

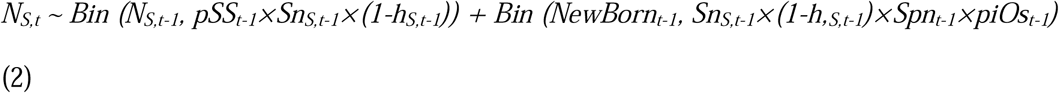

where *Sn*_*j*_ is the survival probability during the first part of the year without hunting, and *1-h*_*j*_ the proportion of females that survived from hunting in the body mass class *j*. Survival probability is defined as *1-NM*_*j*_, with *NM*_*j*_ the natural mortality bringing together all causes of death except hunting (e.g. diseases). The proportion of females shot by hunters in the class *j h*_*j*_ is defined as 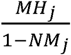, with *MH*_*j*_ the hunting mortality. Assuming that natural mortality was negligible during the hunting period (which is not a very stringent assumption as the multiplicative relationship between natural survival and survival to hunting holds even with some overlap in time between the two sources of mortality), at the end of the year, survival probability for females in the body mass class *j* equals 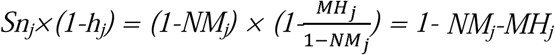 (Lebreton 2005a). *Spn* is the post-natal survival probability (from birth to weaning) and *piOs* the probability for newborn to remain in the small body mass class.

ii. The number of medium females alive in year *t N*_*M,t*_ is the number of small females in year *t-1* that entered the medium class, plus medium females in year *t-1* that remained medium plus the newborn females produced by mature females in year *t-1* that became medium (*1-piOs*) and survived, such as:

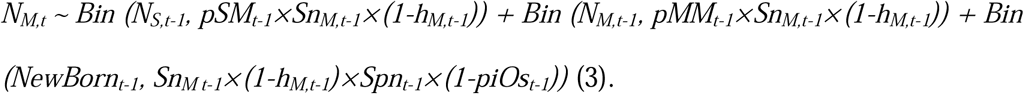

iii. The number of large females alive in year *t N*_*L,t*_ is the number of small and medium females in year *t-1* that entered the large class, plus large females in year *t-1* that survived, such as:

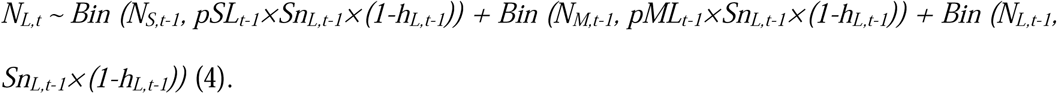

We used a Poisson distribution for the annual number of newborn females produced by small, medium and large females, such as:

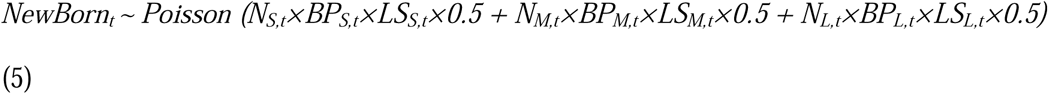

with *BP*_*j,t*_ the annual proportion of breeding females and *LS*_*j,t*_ the annual litter size of each body mass class *j*. We assumed a balanced sex ratio at birth (Servanty et al. 2007).

In addition to the number of females alive in each body mass class, we estimated the annual body mass-specific numbers of females that were shot *NH*_*S,t*_, *NH*_*M,t*_ and *NH*_*L,t*_:

i. The number of small females shot by hunters in year *t NH*_*S,t*_ is the number of small females in year *t-1* that remained small and survived from natural causes but were shot during the hunting season, plus the number of small females produced by mature females in year *t-1* also shot, such as:

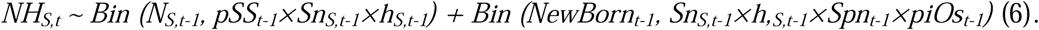
ii. Similarly, the number of medium females shot by hunters in year *t NH*_*M,t*_ is:

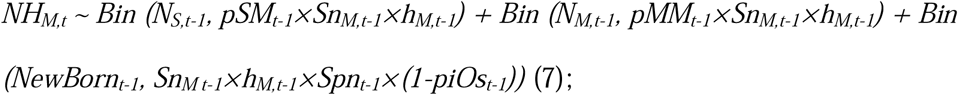
iii. The number of large females shot by hunters in year *t NH*_*L,t*_ is:

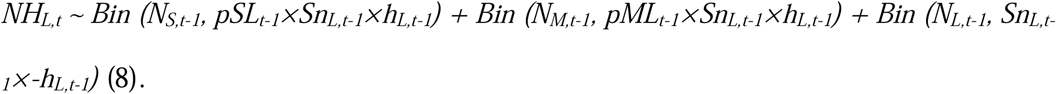

### 2.3 Model for hunting data

Hunting data consisted of the annual numbers of females shot by hunters by body mass classes (small, medium and large), denoted as *y*_*S,t*_, *y*_*M,t*_ and *y*_*L,t*_, respectively. For these data, we used the likelihood of a state-space model (de Valpine & Hastings 2002), which consists of a process model (i.e. the population model previously described, that is the population size as a function of demographic rates and the population size the preceding year) and an observation model. The observation model describes the link between the hunting data *y*_*S,t*_, *y*_*M,t*_ and *y*_*L,t*_ and the true number of females shot by hunters in the population (*NH*_*S,t*_, *NH*_*M,t*_ and *NH*_*L,t*_) (yellow part, Fig. 1). For each body mass class *j*, we assumed that: *y*_*j,t*_ ∼*Normal(NH*_*j,t*_, τ^*j,t*^) truncated to positive values, with 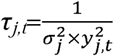, where 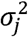 is the term we are estimating. τ^*j,t*^ is the observation error and incorporates both errors in hunting data and lack of fit of the state equations to the true dynamics of the population (see p. 230 in Schaub & Abadi 2011). Note the trick often used in JAGS that consists in defining the distributions by their precision τ^*j,t*^ rather than their variance 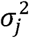, the precision being the inverse of the variance. The likelihood of the state-space model is then the product of the likelihood of the process and the observation equations. The state-space likelihood for hunting data already includes all parameters we want to estimate but they are hardly estimable based on hunting data alone and require additional information coming from CMRR and reproduction data.

### 2.4 Model for capture-mark-recapture-recovery data

CMRR data were analysed using a multistate model (see Lebreton *et al.* 2009 for a review) that allows natural mortality *NM* to be estimated separately from hunting mortality *MH* (Lebreton, Almeras & Pradel 1999; Gamelon *et al.* 2012) for each body mass class (grey part, Fig. 1). We described the fate of an individual using ten states (see Appendix S1). States 1, 2 and 3 were for individuals alive in the small, medium and large body mass classes, respectively. States 4, 5 and 6 were for individuals just shot, again in the three body mass classes, respectively. States 7, 8 and 9 were for individuals that recently died from natural causes, again in the three body mass classes, respectively. The state “dead from a natural cause” cannot be observed because no information was available for individuals that did not die from hunting. State 10 corresponded to individuals already dead. The state “already dead” cannot be observed either but brought together all the dead individuals. To account for the dependency between states (i.e. from year *t* to *t+1*, individuals either survive, die shot by hunters or die from natural causes, see Appendix S1), we used a Dirichlet distribution to model the transitions between states thus ensuring that the sum of these probabilities always equals to 1. The parameters in the multistate model were annual natural mortality *NM*_*j,t*_ and hunting mortality *MH*_*j,t*_ for each body mass class *j*. Moreover, yearly transition probabilities from one body mass class to the next (i.e. *pSS*_*t*_, *pSM*_*t*_, *pSL*_*t*_, *pMM*_*t*_, *pML*_*t*_) were estimated. To account for the dependency between transition probabilities (i.e. *pSL*_*t*_*=1-pSS*_*t*_*-pSM*_*t*_), we used a Dirichlet distribution to model the transitions from the small body mass class to the others. This ensures that the sum of the probabilities (i.e. *pSS*_*t*_ + *pSM*_*t*_ + *pSL*_*t*_) always equals to 1. Regarding the observation process, if an individual was alive, it could be recaptured with probability *p* or not recaptured with probability *1-p*; if an individual just died from hunting, its death could be reported “dead recovery” with probability *r* or not reported with probability *1-r*. Recapture and recovery probabilities were also time and body mass-class dependent (see Appendix S2 and S3).

### 2.5 Model for reproduction data

Reproduction data consisted of the annual numbers of pregnant females shot by body mass classes (small, medium and large), denoted as *R*_*S,t*_, *R*_*M,t*_ and *R*_*L,t*_, respectively and the annual numbers of fetuses counted by body mass classes, denoted as *J*_*S,t*_, *J*_*M,t*_ and *J*_*L,t*_, respectively. We assumed that the number of fetuses per body mass class *J*_*j,t*_ is Poisson distributed such as: *J*_*j,t*_ ∼ *Poisson (R*_*j,t*_*×LS*_*j,t*_), where *LS*_*j,t*_ is the term we are estimating, that is the litter size for a female belonging to the body mass class *j* (blue part, Fig. 1).

### 2.6 Model implementation

Assuming independence among the datasets, the likelihood of the IPM was the product of the likelihoods of the three different datasets (Besbeas *et al.* 2002; Kéry & Schaub 2012): hunting data, CMRR data and reproduction data. The IPM was fit within the Bayesian framework and we used non-informative priors for all the parameters. We only constrained σ^*j*^ in the state-space model to be small using a Uniform distribution between 0 and 0.1, so that the observation errors 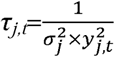 can be large. Markov chain Monte Carlo (MCMC) simulation was used for parameter estimation. All the parameters described in Fig. 1 were estimated within the IPM, even parameters difficult to estimate and often assessed from expert opinion such as *Spn* and *piOs* (see Gamelon *et al.* 2012). The analyses were implemented using JAGS (Plummer 2003) version 4.3.0 called from R version 3.4.3 (R Development Core Team 2017) with package rjags (Plummer 2016). The JAGS code for fitting the IPM is available in Appendix S3.

To assess convergence, we ran three independent chains for 230,000 MCMC iterations, with an adaptation of 180,000 iterations thinning every 100th observation resulting in 1,500 posterior samples. We used the Brooks and Gelman diagnostic R□ to assess the convergence of the simulations and used the rule R□ *<*1.2 to determine whether convergence has been reached (Brooks & Gelman 1998). Convergence was reached for most of the nodes except for four years (1996-1999) in which not all parameters had converged. To ensure that the priors for initial population sizes do not influence estimates of demographic rates and body mass-specific numbers the first year of the study (i.e., in 1991), only the years between 1992 and 2016 were included in the analyses.

## 3 RESULTS

### 3.1 Recapture, recovery and mortality probabilities

Estimated recapture probabilities fluctuated a lot through years, from 0.04 to 0.74 for small females (mean *p*_*S*_=0.36), from 0.03 to 0.82 for medium females (mean *p*_*M*_=0.38) and from 0.002 to 0.62 for large females (mean *p*_*L*_=0.10). These high fluctuations of the average recapture probabilities over years are consistent with earlier studies (see Appendix C in Servanty *et al.* 2010). In accordance with previous work (Toïgo *et al.* 2008; Servanty *et al.* 2010; Gamelon *et al.* 2011), dead recovery reporting probabilities were high for all body mass classes, ranging from 0.75 to 0.97 for small females (mean *r*_*S*_=0.90), from 0.79 to 0.96 for females in the medium body mass class (mean *r*_*M*_=0.88) and from 0.74 to 0.93 for large females (mean *r*_*L*_=0.84). Mortality probability also fluctuated over the study period (Fig. 2 A,B). As expected for this heavily hunted population, hunting mortality *MH* was much higher than natural mortality *NM* for small and medium females. Interestingly, the estimate of the mean post-natal survival was 0.60 [min=0.22; max=0.88] over the study period, and thus close to the value generally set by expert opinion (0.75, see e.g. Gamelon *et al.* 2012).

**Figure 2.**
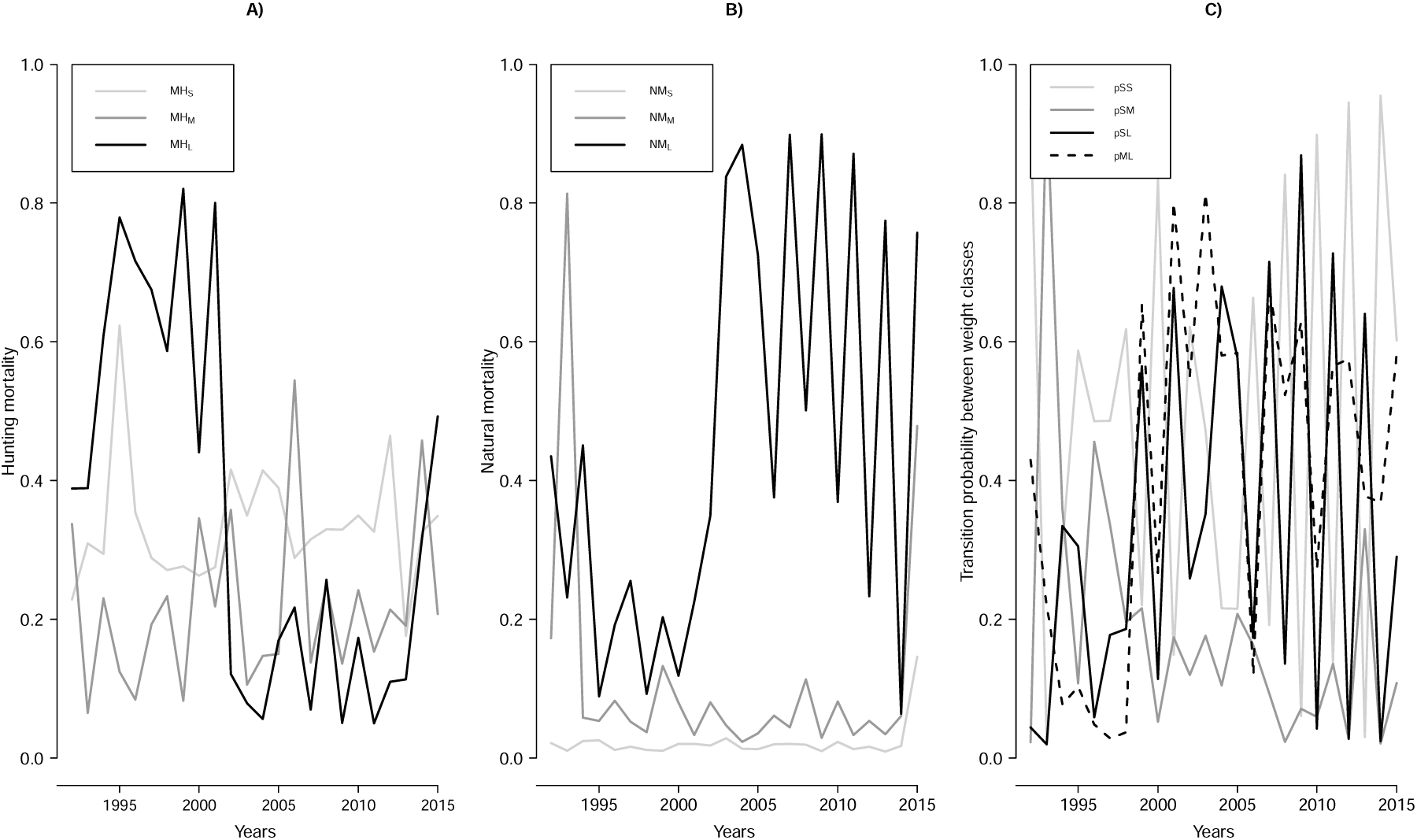
Mortality and growth parameters estimated with the IPM (grey part in Fig. 1). Posterior means of A) annual hunting mortality *MH* and B) annual natural mortality *NM* for each body mass class (small S, medium M and large L). Posterior means of C) transition probability between body mass classes (*pSS*: probability of small females remaining in this class; *pSM*: probability of small females entering the medium class; *pSL*: probability of small females entering the large class; *pML*: probability of medium females entering the large class during the year) for the wild boar population at Châteauvillain-Arc-en-Barrois between 1992 and 2015. CRI are not shown to improve the readability.

### 3.2 Transition probabilities among body-mass classes

Females in the small body mass class had an estimated probability of 0.48 to remain in this class on the average (*pSS*) (Fig. 2C). Alternatively, they moved to the medium-sized class with an estimated probability *pSM* equal to 0.19 on the average, and to the large-sized class with a probability *pSL* of 0.33. Females in the medium body mass class had an estimated probability of 0.41 to enter the large class on the average (*pML*), other medium females remaining in the medium-sized class (*pMM*). Newborn females had an estimated probability of 0.88 [min=0.56; max=0.98] to remain in the small body mass class (*piOs*) on the average, this probability being usually set to 0.60 by expert opinion (see e.g. Gamelon *et al.* 2012).

### 3.3 Reproduction

For reproduction parameters, litter sizes *LS* were strongly body mass-specific, being the largest for large females (mean *LS*_*L*_=6.08 young) and the smallest for females of the small-sized class (mean *LS*_*S*_=4.21 young) (Fig. 3). These results are in accordance with previous studies (Gamelon *et al.* 2013). As for mortality probabilities, litter sizes fluctuated a lot over the study period, from 1.79 to 7.69 young produced for small females (*LS*_*S*_), from 3.91 to 7.50 young for females in the medium weight class (*LS*_*M*_) and from 4.97 to 7 young for large females (*LS*_*L*_). Breeding proportions, i.e. the proportion of breeding females among those in each body mass class, fluctuated over years, especially for females in the small body mass class. These probabilities ranged from 0.36 to 0.84 (mean *BP*_*S*_=0.58) for small females, from 0.43 to 0.65 (mean *BP*_*M*_=0.53) for medium females, and from 0.48 to 0.61 (mean *BP*_*L*_=0.53) for large females.

**Figure 3.**
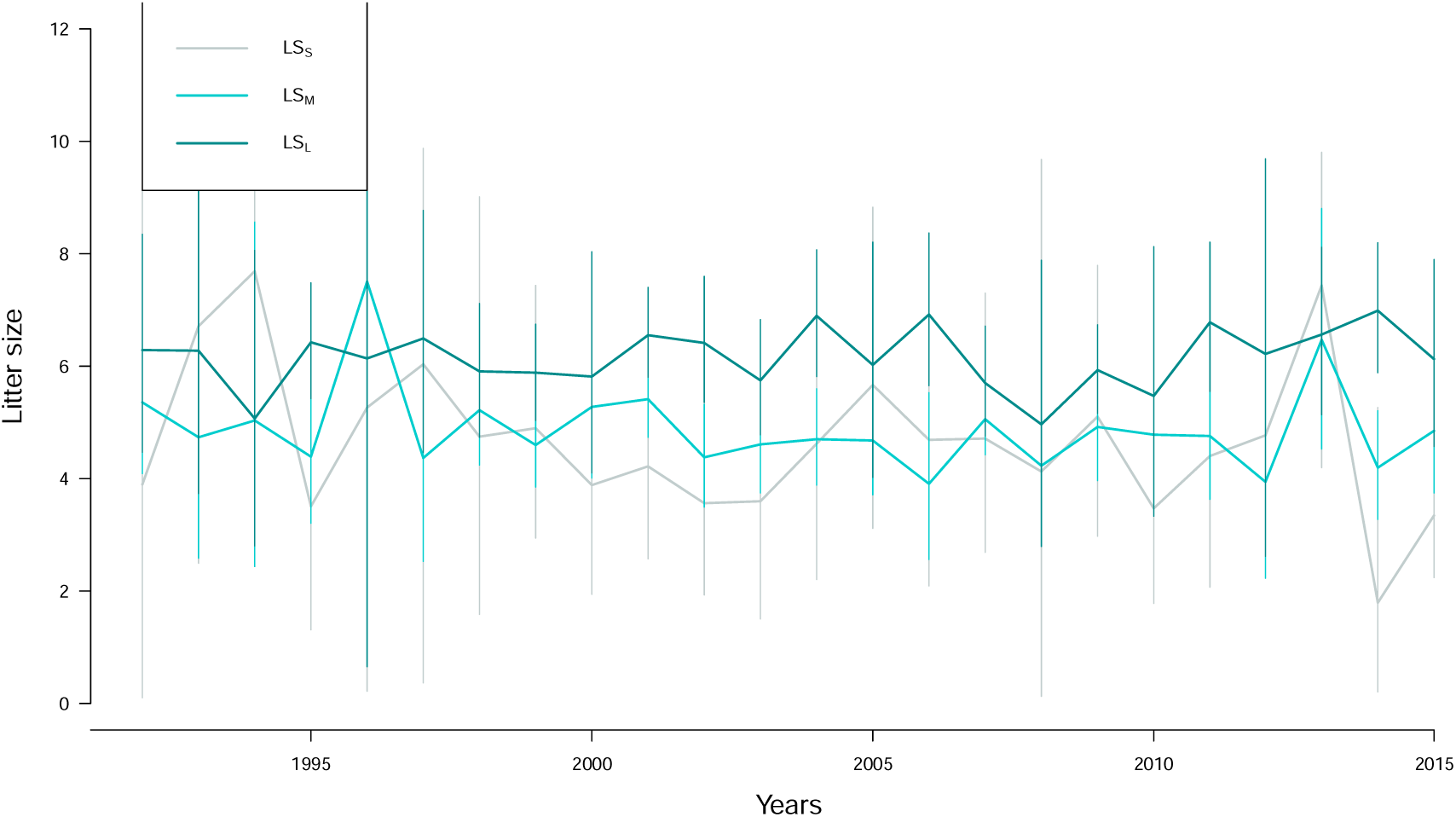
Reproductive parameters estimated with the IPM (blue part in Fig. 1). Posterior means of annual litter size *LS* for each body mass class (small S, medium M and large L) (together with their 95% CRI) for the wild boar population at Châteauvillain-Arc-en-Barrois between 1992 and 2015.

### 3.4 Population sizes

High annual fluctuations in mortality, transition between body mass classes and reproductive rates translated to high variation in population size over years (Fig. 4). Each year, with a mean of *NH*_*S*_=143 females, most of the females removed from hunting belonged to the small-sized class (Fig. 4A). The mean number of females shot by hunters in the medium class *NH*_*M*_ was estimated to be 81, whereas they were very few large females in the hunting bags (mean *NH*_*L*_=35). Noticeably, the numbers of females shot in each body mass class expected from the IPM was very close to the observed numbers *y*_*j*_ (dots in Fig. 4A). Observation errors σ were estimated at 0.096 for small and medium females and 0.099 for large females. In terms of numbers of females alive in the population, small females constituted the largest part of the population (mean number of small females *N*_*S*_=290), followed by medium (mean *N*_*M*_=77) and then large females (mean *N*_*L*_=36) (Fig. 4B).

**Figure 4.**
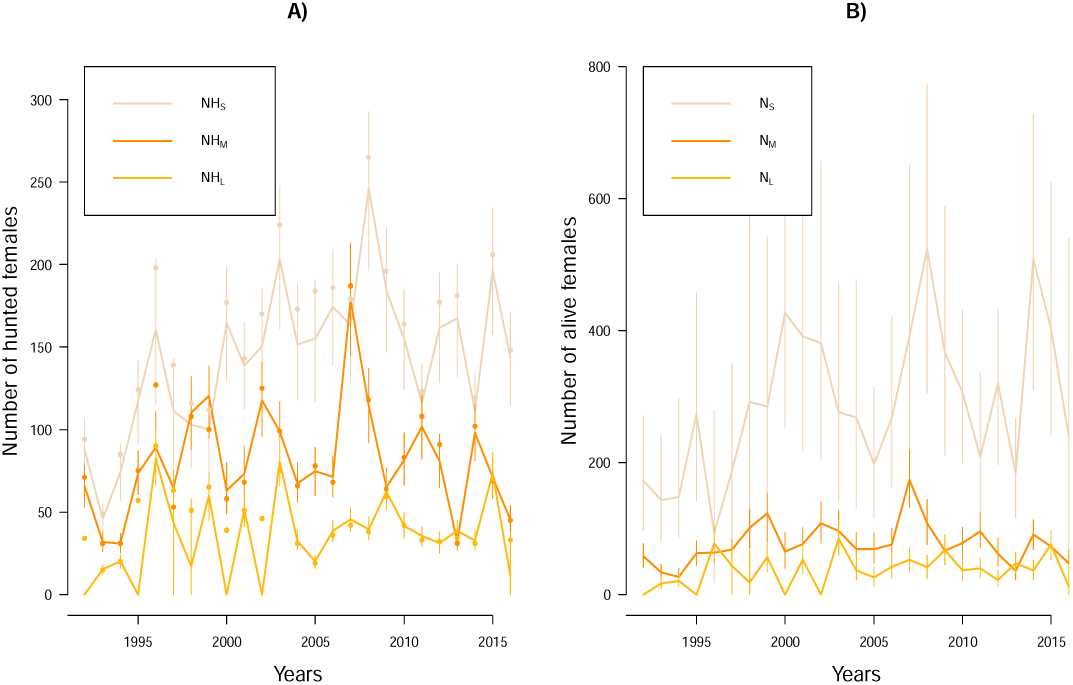
Population sizes estimated with the IPM (yellow part in Fig. 1). Posterior means (together with their 95% CRI) of A) annual numbers of females shot by hunters *NH* and B) annual numbers of females alive *N* in each body mass class (small S, medium M and large L) for the wild boar population at Châteauvillain-Arc-en-Barrois between 1992 and 2016. Dots correspond to observed numbers of females shot by hunters in the three body mass classes (*y*_*S*_, *y*_*M*_ and *y*_*L*_).

## 4 DISCUSSION

We develop here an IPM that makes efficient use of data commonly collected in exploited populations, i.e. body mass at captures/death as well as number of individuals removed by harvesting. Our model makes it possible to directly compare the observed and the expected number of individuals shot. It also provides accurate and precise estimates of key demographic parameters, including some that cannot be estimated from separate analyses. Using a wild boar population as a case study, we demonstrate that this framework is a powerful tool to gain a good understanding of the dynamics of exploited populations.

### 4.1 A comprehensive picture of population dynamics

In accordance with previous work (Toïgo *et al.* 2008; Gamelon *et al.* 2011), we showed that hunting mortality probabilities *MH* are high for all body mass classes. Conversely, natural mortality *NM* was low (Fig. 2 A,B). This is expected among ungulates where the average natural adult survival probability *Sn* often exceed 0.95 in females (Gaillard *et al.* 2000). These hunting and natural mortality patterns are in accordance with other hunted wild boar populations (see Gamelon 2019 for a review). Our IPM thus confirms that hunting is an important driver of wild boar population dynamics. Regarding to reproduction parameters, wild boar is a highly fecund species, being able to produce large litters (Fig. 3) as early as their first year of life (Servanty *et al.* 2009) at a body mass lower than 30 kg (small mass-class). Together with reduced survival due to hunting, this leads to a short generation time, i.e. a low mean age of mothers at childbirth (Gaillard *et al.* 2005, 2016) close to two years, whereas it is close to six years for similar-sized ungulates (Servanty et al. 2011). This unusual life history among ungulates (Focardi *et al.* 2008), characterized with a fast turnover of individuals in the population, explains why the number of individuals in the population did not collapse (Fig. 4B) during the study period despite such a high hunting pressure, and why younger/lighter females constitute more than 50% of the individuals alive. By analyzing simultaneously CMRR data, reproductive data and hunting data, and by estimating jointly all the demographic parameters, our IPM offers a comprehensive picture of the underlying demographic mechanisms allowing this population not to collapse over the last 25 years despite a high hunting pressure.

### 4.2 Estimates for demographic parameters hardly measurable in the field

The size of exploited populations is generally poorly known, and the IPM allows us to get accurate and precise estimates even in the absence of surveys of the number of individuals alive, which are usually needed to make relevant management recommendations. Interestingly, the model also allows us to estimate some growth parameters that are difficult to measure in the field, because they would require multiple captures of the same individuals over their lifespan. We found marked year-to-year variation in the probability for a female to remain in the same body mass class or enter a heavier body mass class (Fig. 2C). A large body of empirical evidence shows that increasing density is generally associated with reduced body mass in large herbivores (Bonenfant *et al.* 2009). Fluctuations in the strength of density dependence in body growth could explain such among-year variation in transition probabilities. Factors affecting transitions between body mass classes remain to be carefully explored, offering exciting avenues of research. Likewise, the IPM allowed us to get annual estimates of post-natal survival, a parameter often tricky to estimate empirically. For instance, Baubet et al. (2009) aimed to tag piglets inside their birth nest to assess post-natal survival from birth to weaning. They failed in this task not only because of expected difficulties to locate the birth nests, but also because it induced abandonment of the piglets after tagging. By jointly analysing different data sources, the IPM is a powerful tool to achieve such a goal.

### 4.3 A framework based on data commonly collected in exploited populations

To make harvest sustainable, i.e. avoid overharvest of declining populations or avoid applying too low harvest rate to increasing populations, the dynamics of exploited populations should be fully understood. In frequent situations where data are limited, it might be a tricky task. Our IPM is not solely a powerful tool to understand the dynamics of increasing wild boar populations, but it is clearly applicable to many other populations in both terrestrial and marine environments. For instance, a lot of commercially important marine fish species are subject to strong harvesting (Pauly *et al.* 2002; Hutchings & Reynolds 2004). The IPM can be a suitable tool to anticipate the collapse of some of them (Hutchings & Myers 1994; Myers, Hutchings & Barrowman 1997) by making efficient use of the limited data available (see Maunder 2004; Saunders, Cuthbert & Zipkin 2018 for the use of IPM in prospective analyses).

The use of IPM as a management tool is not novel in fisheries (see Maunder & Punt 2013 for a review) but they are to date based on age-structured population models. However, fish are indeterminate growers for which demographic parameters are usually strongly mass-dependent. Moreover, while taking reliable mass measurements of animals killed by humans is straightforward, information on age is more cumbersome and often inaccurate and thus recorded in very few populations only. Even in determinate growers such as birds and mammals, evidence is accumulating that body mass is a crucial structuring factor of population dynamics (Gaillard *et al.* 2000 for a review on large herbivores; see Coulson, Tuljapurkar & Childs 2010 for a case study on Soay sheep *Ovis aries*), making biologically relevant the use of body mass-structured models in IPM for a large range of exploited populations of vertebrates. More generally, our framework based on a body mass-structured model adds to the spate of studies that have recently flourished in the literature showing that trait-based approaches (such as body mass) and demographic approaches are intertwined (Salguero-Gómez *et al.* 2018; Smallegange & Berg 2019). Another clear advantage of our IPM is the inclusion of the number of dead individuals in the body mass-structured population model, whereas all the IPMs built so far only included the number of individuals alive in a population model (Caswell 2001). In exploited populations, getting information on the number of individuals alive is challenging whereas information on the number of individuals killed (by hunting or fishing) is often available. This number is even almost perfectly known in our population. With the inclusion of information related to hunting bags, our IPM renders possible a direct comparison between observed and predicted numbers of individuals shot in each body-mass class, a crucial demographic information in exploited populations.

### 4.4 Possible extensions

The use of the number of individuals harvested in our IPM could be modified to the use of proportions. A classical case is that of duck hunting, in which programs for collecting samples of wings of individuals shot make it possible to estimate the proportion of first-year and after first-year (i.e. adult) individuals of both sexes in the harvest (Raftovich, Chandler & Fleming 2018). Such estimates of age-and sex structures among dead individuals have been used up to now in an ad hoc fashion to match model predictions (Holopainen *et al.* 2018). The analysis of such data based on an IPM would certainly greatly improve our knowledge and understanding of the differences in vulnerability to hunting between young and adults and between sexes in duck populations.

Another possible straightforward extension of our IPM could be the modification of the number and size of the body mass classes. Beyond body mass, the IPM is generalizable to all kinds of states, such as breeder/non breeder which is a commonly reported state in CMR studies on birds (see e.g. Pardo, Barbraud & Weimerskirch 2014 for a multistate model on wandering albatross *Diomedea exulans*). Beyond mortality due to hunting and fishing, the IPM can also be expanded to all forms of anthropogenic mortalities (see e.g. Chevallier *et al.* 2015 for mortality induced by electrocution in Bonelli’s eagle *Aquila fasciata*). Finally, while we used non-informative priors and allowed all the parameters to be estimated within the IPM, it is straightforward to integrate a priori biological knowledge by setting some parameters to known values or by using informative priors.

## Conclusions

The recent methodological advances, such as the introduction of MCMC that have highly contributed to the expansion of IPM, allowed us to propose an IPM that makes efficient use of (limited) harvest data for the monitoring of exploited populations. The model is flexible and can be adapted to the life history and the data available for the population of interest. This model that integrates commonly collected data in marine and terrestrial exploited populations is therefore widely applicable.

## ACKNOWLEDGMENTS

Wild boar data collection was performed and granted by the French National Agency for Wildlife and Hunting (ONCFS). We are grateful to all those who helped marking and collecting harvested wild boar, in particular Serge Brandt, Eveline Nivois and Cyril Rousset, to the Office National des Forêts and to F. Jehlé who allowed us to work on the study area. This work was supported by the Research Council of Norway through its Centre of Excellence funding scheme, Project Number 223257.

## AUTHOR’S CONTRIBUTIONS

EB contributed to data collection. MG, EB, AB, JMG, JDL and OG conceived the study. MG conducted the analyses with insights from OG, wrote the first draft and all authors contributed to revisions of the initial manuscript.

## DATA AVAILABILITY

The data supporting the results will be archived in an appropriate public repository in Dryad, should the manuscript be accepted.

