## Supplementary material for "Efficient use of harvest data: An integrated population model for exploited animal populations": Supp material

**Appendix S1**

Matrix showing transitions from state at year *t* (rows) to state at year *t+1* (columns). Ten states are considered: alive, just shot, just died from a natural cause and already dead for small *S*, medium-sized *M* or large *L* body body mass-classes. Individuals shot, dead of a natural cause or already dead at year *t* remain in the state “Dead” (probability=1). See figure 1 for a definition of the parameters, all time-dependent.

$\begin{matrix} & \mathrm{Small} \mathrm{alive} & \mathrm{Medium} \mathrm{alive} & \mathrm{Large} \mathrm{alive} & \mathrm{Small} h\mathrm{unted} & \mathrm{Medium} h\mathrm{unted} & \mathrm{Large} h\mathrm{unted} & \mathrm{Small} \mathrm{nat} & \mathrm{Medium} \mathrm{nat} & \mathrm{Large} \mathrm{nat} & \mathrm{Dead} \\ \mathrm{Small} \mathrm{alive} & \mathrm{pSS}_{t}\timesɸ_{S1,t} & \mathrm{pSM}_{t}\timesɸ_{S1,t} & \mathrm{pSL}_{t}\timesɸ_{S1,t} & ɸ_{S2,t} & ɸ_{S3,t} & ɸ_{S4,t} & ɸ_{S5,t} & ɸ_{S6,t} & ɸ_{S7,t} & 0 \\ \mathrm{Medium} \mathrm{alive} & 0 & \mathrm{pMM}_{t}\timesɸ_{M1,t} & \mathrm{pML}_{t}\timesɸ_{M1,t} & 0 & ɸ_{M2,t} & ɸ_{M3,t} & 0 & ɸ_{M4,t} & ɸ_{M5,t} & 0 \\ \mathrm{Large} \mathrm{alive} & 0 & 0 & ɸ_{L1,t} & 0 & 0 & ɸ_{L2,t} & 0 & 0 & ɸ_{L3,t} & 0 \\ \mathrm{Small} h\mathrm{unted} & 0 & 0 & 0 & 0 & 0 & 0 & 0 & 0 & 0 & 1 \\ \mathrm{Medium} h\mathrm{unted} & 0 & 0 & 0 & 0 & 0 & 0 & 0 & 0 & 0 & 1 \\ \mathrm{Large} h\mathrm{unted} & 0 & 0 & 0 & 0 & 0 & 0 & 0 & 0 & 0 & 1 \\ \mathrm{Small} \mathrm{nat} & 0 & 0 & 0 & 0 & 0 & 0 & 0 & 0 & 0 & 1 \\ \mathrm{Medium} \mathrm{nat} & 0 & 0 & 0 & 0 & 0 & 0 & 0 & 0 & 0 & 1 \\ \mathrm{Large} \mathrm{nat} & 0 & 0 & 0 & 0 & 0 & 0 & 0 & 0 & 0 & 1 \\ \mathrm{Dead} & 0 & 0 & 0 & 0 & 0 & 0 & 0 & 0 & 0 & 1 \end{matrix}$

Natural mortality:

- for small females: NM_s,t_=$ɸ_{S5,t}$;
- for medium females: NM_M,t_ =$ɸ_{S6,t}+ɸ_{M4,t}$;
- for large females: NM_L,t_ =$ɸ_{S7,t}+ɸ_{M5,t}+ɸ_{L3,t}$.

Hunting mortality:

- for small females: MH_s,t_ =$ɸ_{S2,t}$;
- for medium females: MH_M,t_ =$ɸ_{S3,t}+ɸ_{M2,t}$;
- for large females: MH_L,t_ =$ɸ_{S4,t}+ɸ_{M3,t}+ɸ_{L2,t}$.

**Appendix S2**

Matrix showing recapture probability (*p_j,t_*) and recovery probability (*r_j,t_*) at year *t* for body mass class *j*. States are in row and observations in column (0: female not seen, 1, 2, 3: female recaptured in the small, medium and large body mass-class, respectively; 4, 5, 6: female shot by hunters in the small, medium and large body mass-class, respectively). Females not seen (0) but alive are not captured (*1-p_j,t_*); females alive and seen (1) are captured (*p_j,t_*); females shot and not seen (0) are not recovered (*1-r_j,t_*); females shot are recovered with a recovery probability (*r_j,t_*). Females dead from a natural cause are never seen (0), same for dead females (probability=1).

$$\begin{matrix} & 0 & 1 & 2 & 3 & 4 & 5 & 6 \\ Small alive & 1-p_{S,t} & p_{S,t} & 0 & 0 & 0 & 0 & 0 \\ Medium alive & 1-p_{M,t} & 0 & p_{M,t} & 0 & 0 & 0 & 0 \\ Large alive & 1-p_{L,t} & 0 & 0 & p_{L,t} & 0 & 0 & 0 \\ Small hunted & 1-r_{S,t} & 0 & 0 & 0 & r_{S,t} & 0 & 0 \\ Medium hunted & 1-r_{M,t} & 0 & 0 & 0 & 0 & r_{M,t} & 0 \\ Large hunted & 1-r_{L,t} & 0 & 0 & 0 & 0 & 0 & r_{L,t} \\ Small nat & 1 & 0 & 0 & 0 & 0 & 0 & 0 \\ Medium nat & 1 & 0 & 0 & 0 & 0 & 0 & 0 \\ Large nat & 1 & 0 & 0 & 0 & 0 & 0 & 0 \\ \mathrm{Dead} & 1 & 0 & 0 & 0 & 0 & 0 & 0 \end{matrix}$$

**Appendix S3**

Code JAGS used to implement the integrated population model.

##################################################################

\# 0 - READ IN DATA

##################################################################

setwd("")

mydata <- as.matrix(mydata) #Load CMRR data

J=as.matrix(J) # Load number of fetuses counted per year (rows) for small, medium and large females (columns)

R=as.matrix(R) # Load number of small, medium and large females (in columns) pregnant annually (in rows)

\# Number of harvested females in the three weight classes (small, medium and large) between 1991-2016:

yHS <- c(166, 94, 49, 85, 124, 198, 139, 116, 112, 177, 143, 170, 224, 173, 184, 186, 179, 265, 196, 164, 123, 177, 181, 119, 206, 148)

yHM <-c(56, 71, 31, 31, 75, 127, 53, 108, 100, 58, 68, 125, 99, 66, 78, 68, 187, 118, 64, 83, 108, 91, 31, 102, 69, 45)

yHL <- c(34, 34, 15, 20, 57, 90, 63, 51, 65, 39, 51, 46, 75, 31, 19, 36, 42, 38, 59, 42, 33, 32, 34, 31, 70, 33)

effHS <- c(166, 94, 49, 85, 124, 198, 139, 116, 112, 177, 143, 170, 224, 173, 184, 186, 179, 265, 196, 164, 123, 177, 181, 119, 206, 148)

effHM <-c(56, 71, 31, 31, 75, 127, 53, 108, 100, 58, 68, 125, 99, 66, 78, 68, 187, 118, 64, 83, 108, 91, 31, 102, 69, 45)

effHL <- c(34, 34, 15, 20, 57, 90, 63, 51, 65, 39, 51, 46, 75, 31, 19, 36, 42, 38, 59, 42, 33, 32, 34, 31, 70, 33)

##################################################################

\# 1 - DATA MANIPULATION

##################################################################

\# Data are in matrix “mydata”, one female per row, with coding as follows:

\# 0 = female not observed;

\# 1 = female observed Small;

\# 2 = female observed Medium;

\# 3 = female observed Large;

\# 4 = female found dead Small;

\# 5 = female found dead Medium;

\# 6 = female found dead Large;

\# Number of individuals

n <- dim(mydata)[[1]]

\# Number of capture occasions

K <- dim(mydata)[[2]]

\# Compute date of first capture

e <- NULL

last <- NULL

quid <- NULL

for (i in 1:n){

temp <- 1:K

quid <- c(quid,(mydata[i,min(temp[mydata[i,]>=1])]))

e <- c(e,min(temp[mydata[i,]>=1]))

}

for (i in 1:n){

temp <- 1:K

mask = (mydata[i,]>=4)

if (sum(mask)==1) {last <- c(last,temp[mask])}

else {last <- c(last,K)}

}

\# Number of years

nyears <- length(yHS)

##################################################################

\# 2 – SPECIFY MODEL IN BUGS LANGUAGE

##################################################################

sink("ipm-prod.bug")

cat("

model{

#-------------------------------------------------

\# 1. Define the priors for the parameters

#-------------------------------------------------

\# Initial population sizes

\# Number of small females alive NS[t,1]

fvs ~ dnorm(50,50)T(0,)

NS[1,1] <- round(fvs)

\# Number of medium females alive NM[t,1]

fvm ~ dnorm(50,50)T(0,)

NM[1,1] <- round(fvm)

\# Number of large females alive NL[t,1]

fvl ~ dnorm(50,50)T(0,)

NL[1,1] <- round(fvl)

\# Initial state

piVS ~ dunif(0,1)

for (t in 1:(nyears-1)){

\# Dirichlet prior for survival probabilities

survS[1:7,t] ~ ddirch(alphaS[])

survM[1:5,t] ~ ddirch(alphaM[])

survL[1:3,t] ~ ddirch(alphaL[])

psiML[t] ~ dunif(0,1) # Transition from medium to large-sized body mass class (=pML=1-pMM)

pSMSL[1:3,t] ~ ddirch(transitionS[]) # Transitions from small to small, medium and large- sized body mass classes (=pSS, pSM, pSL)

ppS[t] ~ dunif(0,1) # Recapture probability for small females (=p_S_)

ppM[t] ~ dunif(0,1) # Recapture probability for medium females (=p_M_)

ppL[t] ~ dunif(0,1) # Recapture probability for large females (=p_L_)

llS[t] ~ dunif(0,1) # Recovery probability for small females (=r_S_)

llM[t] ~ dunif(0,1) # Recovery probability for medium females (=r_M_)

llL[t] ~ dunif(0,1) # Recovery probability for large females (=r_L_)

\# Hunting mortality

pS[2,t] <- survS[2,t] # Small (=MH_s_)

pM[2,t] <- survM[2,t]+survS[3,t] # Medium (=MH_M_)

pL[2,t] <- survS[4,t]+survM[3,t]+survL[2,t] # Large (=MH_L_)

\# Natural mortality

pS[3,t] <- survS[5,t] # Small (=NM_s_)

pM[3,t] <- survS[6,t]+survM[4,t] # Medium (=NM_M_)

pL[3,t] <- survS[7,t]+survM[5,t]+survL[3,t] # Large (=NM_L_)

\# Natural survival = 1-Natural mortality

pS[1,t] <- 1-pS[3,t]

pM[1,t] <- 1-pM[3,t]

pL[1,t] <- 1-pL[3,t]

\# Proportion of hunted individuals h = Hunting mortality/Natural survival

h[1,t] <- pS[2,t]/pS[1,t] # Small

h[2,t] <- pM[2,t]/pM[1,t] # Medium

h[3,t] <- pL[2,t]/pL[1,t] # Large

\# Post-natal survival

SPN[t] ~ dunif(0,1)

\# Probability for newborn to remain in the small weight class (piOs)

piOS[t] ~ dunif(0,1)

\# Breeding proportions

BPS[t] ~ dunif(0,1) # Small

BPM[t] ~ dunif(0,1) # Medium

BPL[t] ~ dunif(0,1) # Large

\# Litter size (i.e. number of newborn produced)

FECUND_S[t]~ dunif(0,10) # Small (=LS_s_)

FECUND_M[t]~ dunif(0,10) # Medium (=LS_M_)

FECUND_L[t]~ dunif(0,10) # Large (=LS_L_)

}

#-------------------------------------------------

\# 2. The likelihoods of the single data sets

#-------------------------------------------------

#------------------------------------------------

\# 2.1. Likelihood for population data

#------------------------------------------------

\# 2.1.1 System process

for (t in 2:nyears){

\# Reproduction (with balanced sex-ratio at birth)

fecS[t-1] <- FECUND_S[t-1]*0.5*BPS[t-1]*NS[t-1,1] # Small

fecM[t-1] <- FECUND_M[t-1]*0.5*BPM[t-1]*NM[t-1,1] # Medium

fecL[t-1] <- FECUND_L[t-1]*0.5*BPL[t-1]*NL[t-1,1] # Large

\# Round all population sizes

NSr[t-1,1]<-round(NS[t-1,1])

fec_r_p[t-1]~ dpois(fecM[t-1]+fecL[t-1]+fecS[t-1]) (=newborn)

fec_r[t-1]<-round(fec_r_p[t-1])

NMr[t-1,1]<-round(NM[t-1,1])

NLr[t-1,1]<-round(NL[t-1,1])

\# Number of individuals alive in each body mass class

\# Number of small females = females that remained in this class and survived + females produced by small, medium and large females

NSS[t]~ dbin(pS[1,t-1]*(1-h[1,t-1])*pSMSL[1,t-1],NSr[t-1,1]) # Small remaining small and surviving

NSfromML[t]~ dbin(pS[1,t-1]*(1-h[1,t-1])*SPN[t-1]*piOS[t-1],fec_r[t-1]) # Small produced by small, medium & large females that survived and remained in this class

NS[t,1]<-NSS[t]+NSfromML[t]

\# Number of medium females = small females that moved to the medium class within the year and survived + medium females that remained in this class and survived + females produced by small, medium & large females that moved to the medium class

NSM[t]~ dbin(pM[1,t-1]*ifelse (1-h[2,t-1]<0, 1-h[2,t-1]==0, 1-h[2,t-1]==1-h[2,t-1])*pSMSL[2,t-1],NSr[t-1,1]) # Small becoming medium and surviving

NMM[t]~ dbin(pM[1,t-1]*ifelse (1-h[2,t-1]<0, 1-h[2,t-1]==0, 1-h[2,t-1]==1-h[2,t-1])*(1-psiML[t-1]),NMr[t-1,1]) # Medium remaining medium and surviving

NMfromML[t]~ dbin(pM[1,t-1]*ifelse (1-h[2,t-1]<0, 1-h[2,t-1]==0, 1-h[2,t-1]==1-h[2,t-1])*SPN[t-1]*(1-piOS[t-1]),fec_r[t-1]) # Small produced by small, medium & large females that survived and moved to the medium class

NM[t,1]<-NSM[t]+NMM[t]+NMfromML[t]

\# Number of large females = large females already in this class that survived + medium females that survived and entered the large class + small females that survived and entered the large class

NSL[t]~ dbin(pL[1,t-1]*ifelse (1-h[3,t-1]<0, 1-h[3,t-1]==0, 1-h[3,t-1]==1-h[3,t-1])*pSMSL[3,t-1],NSr[t-1,1]) # Small becoming large and surviving

NLfromL[t]~ dbin(pL[1,t-1]*ifelse (1-h[3,t-1]<0, 1-h[3,t-1]==0, 1-h[3,t-1]==1-h[3,t-1]),NLr[t-1,1]) # Large remaining large and surviving

NLfromM[t]~ dbin(pL[1,t-1]*ifelse (1-h[3,t-1]<0, 1-h[3,t-1]==0, 1-h[3,t-1]==1-h[3,t-1])*psiML[t-1],NMr[t-1,1]) # Medium becoming large and surviving

NL[t,1]<-NSL[t]+NLfromL[t]+NLfromM[t]

\## Number of individuals killed by hunting in each body mass class

\# Number of small females = females that remained in this class and died from hunting + females produced by small, medium & large females and died from hunting

NSSh[t]~ dbin(pS[1,t-1]*h[1,t-1]*pSMSL[1,t-1],NSr[t-1,1]) # Small remaining small and being killed by hunting

NSfromMLh[t]~ dbin(pS[1,t-1]*h[1,t-1]*SPN[t-1]*piOS[t-1],fec_r[t-1]) # Small produced by small, medium & large females that remained in this class and died from hunting

NS[t,2]<-NSSh[t]+NSfromMLh[t]

\# Number of medium females= small females that moved to the medium class and died from hunting + medium females that remained in this class and died from hunting + females produced by small, medium & large females that moved to the medium class and died from hunting

NSMh[t]~ dbin(pM[1,t-1]*ifelse (h[2,t-1]>1, h[2,t-1]==1, h[2,t-1]==h[2,t-1])*pSMSL[2,t-1],NSr[t-1,1]) # Small becoming medium and being killed by hunting

NMMh[t]~ dbin(pM[1,t-1]*ifelse (h[2,t-1]>1, h[2,t-1]==1, h[2,t-1]==h[2,t-1])*(1-psiML[t-1]),NMr[t-1,1]) # Medium remaining medium and killed by hunting

NMfromMLh[t]~ dbin(pM[1,t-1]*ifelse (h[2,t-1]>1, h[2,t-1]==1, h[2,t-1]==h[2,t-1])*SPN[t-1]*(1-piOS[t-1]),fec_r[t-1]) # Small produced by small, medium & large females that moved to the medium class and were killed by hunting

NM[t,2]<-NSMh[t]+NMMh[t]+NMfromMLh[t]

\# Number of large females = large females already in this class that died from hunting + medium females that died from hunting and entered the large class + small females that entered the large class and died from hunting

NSLh[t]~ dbin(pL[1,t-1]*ifelse (h[3,t-1]>1, h[3,t-1]==1, h[3,t-1]==h[3,t-1])*pSMSL[3,t-1],NSr[t-1,1]) # Small becoming large and killed by hunting

NLfromLh[t]~ dbin(pL[1,t-1]*ifelse (h[3,t-1]>1, h[3,t-1]==1, h[3,t-1]==h[3,t-1]),NLr[t-1,1]) # Large remaining large and killed by hunting

NLfromMh[t]~ dbin(pL[1,t-1]*ifelse (h[3,t-1]>1, h[3,t-1]==1, h[3,t-1]==h[3,t-1])*psiML[t-1],NMr[t-1,1]) # Medium becoming large and killed by hunting

NL[t,2]<-NSLh[t]+NLfromLh[t]+NLfromMh[t]

}

\# 2.1.2 Observation process

\# Observation error / coefficient of variation

sigyS ~ dunif(0,0.1)

sigyM ~ dunif(0,0.1)

sigyL ~ dunif(0,0.1)

for (t in 2:nyears){

varyS[t] <- sigyS*sigyS*effHS[t]*effHS[t]

varyM[t] <- sigyM*sigyM*effHM[t]*effHM[t]

varyL[t] <- sigyL*sigyL*effHL[t]*effHL[t]

tauyS[t] <- 1/varyS[t]

tauyM[t] <- 1/varyM[t]

tauyL[t] <- 1/varyL[t]

yHS[t] ~ dnorm(NS[t,2], tauyS[t])T(0,)

yHM[t] ~ dnorm(NM[t,2], tauyM[t])T(0,)

yHL[t] ~ dnorm(NL[t,2], tauyL[t])T(0,)

}

#----------------------------------------------------------------

\# 2.2 Likelihood for capture-recapture-recovery data

#----------------------------------------------------------------

\# Probabilities for each initial state

px0[1] <- piVS # Probability of being in initial state alive small

px0[2] <- 1 - piVS # Probability of being in initial state alive medium

px0[3] <- 0

px0[4] <- 0

px0[5] <- 0

px0[6] <- 0

px0[7] <- 0

px0[8] <- 0

px0[9] <- 0

px0[10] <- 0

po.init[1,1] <- 0

po.init[1,2] <- 1

po.init[1,3] <- 0

po.init[1,4] <- 0

po.init[1,5] <- 0

po.init[1,6] <- 0

po.init[1,7] <- 0

po.init[2,1] <- 0

po.init[2,2] <- 0

po.init[2,3] <- 1

po.init[2,4] <- 0

po.init[2,5] <- 0

po.init[2,6] <- 0

po.init[2,7] <- 0

po.init[3,1] <- 0

po.init[3,2] <- 0

po.init[3,3] <- 0

po.init[3,4] <- 1

po.init[3,5] <- 0

po.init[3,6] <- 0

po.init[3,7] <- 0

po.init[4,1] <- 0

po.init[4,2] <- 0

po.init[4,3] <- 0

po.init[4,4] <- 0

po.init[4,5] <- 1

po.init[4,6] <- 0

po.init[4,7] <- 0

po.init[5,1] <- 0

po.init[5,2] <- 0

po.init[5,3] <- 0

po.init[5,4] <- 0

po.init[5,5] <- 0

po.init[5,6] <- 1

po.init[5,7] <- 0

po.init[6,1] <- 0

po.init[6,2] <- 0

po.init[6,3] <- 0

po.init[6,4] <- 0

po.init[6,5] <- 0

po.init[6,6] <- 0

po.init[6,7] <- 1

po.init[7,1] <- 1

po.init[7,2] <- 0

po.init[7,3] <- 0

po.init[7,4] <- 0

po.init[7,5] <- 0

po.init[7,6] <- 0

po.init[7,7] <- 0

po.init[8,1] <- 1

po.init[8,2] <- 0

po.init[8,3] <- 0

po.init[8,4] <- 0

po.init[8,5] <- 0

po.init[8,6] <- 0

po.init[8,7] <- 0

po.init[9,1] <- 1

po.init[9,2] <- 0

po.init[9,3] <- 0

po.init[9,4] <- 0

po.init[9,5] <- 0

po.init[9,6] <- 0

po.init[9,7] <- 0

po.init[10,1] <- 1

po.init[10,2] <- 0

po.init[10,3] <- 0

po.init[10,4] <- 0

po.init[10,5] <- 0

po.init[10,6] <- 0

po.init[10,7] <- 0

for (t in 1:(nyears-1)){

\# Probabilities of observations at a given occasion given states at this occasion (see Appendix S2)

po[1,t,1] <- 1-ppS[t]

po[1,t,2] <- ppS[t]

po[1,t,3] <- 0

po[1,t,4] <- 0

po[1,t,5] <- 0

po[1,t,6] <- 0

po[1,t,7] <- 0

po[2,t,1] <- 1-ppM[t]

po[2,t,2] <- 0

po[2,t,3] <- ppM[t]

po[2,t,4] <- 0

po[2,t,5] <- 0

po[2,t,6] <- 0

po[2,t,7] <- 0

po[3,t,1] <- 1-ppL[t]

po[3,t,2] <- 0

po[3,t,3] <- 0

po[3,t,4] <- ppL[t]

po[3,t,5] <- 0

po[3,t,6] <- 0

po[3,t,7] <- 0

po[4,t,1] <- 1-llS[t]

po[4,t,2] <- 0

po[4,t,3] <- 0

po[4,t,4] <- 0

po[4,t,5] <- llS[t]

po[4,t,6] <- 0

po[4,t,7] <- 0

po[5,t,1] <- 1-llM[t]

po[5,t,2] <- 0

po[5,t,3] <- 0

po[5,t,4] <- 0

po[5,t,5] <- 0

po[5,t,6] <- llM[t]

po[5,t,7] <- 0

po[6,t,1] <- 1-llL[t]

po[6,t,2] <- 0

po[6,t,3] <- 0

po[6,t,4] <- 0

po[6,t,5] <- 0

po[6,t,6] <- 0

po[6,t,7] <- llL[t]

po[7,t,1] <- 1

po[7,t,2] <- 0

po[7,t,3] <- 0

po[7,t,4] <- 0

po[7,t,5] <- 0

po[7,t,6] <- 0

po[7,t,7] <- 0

po[8,t,1] <- 1

po[8,t,2] <- 0

po[8,t,3] <- 0

po[8,t,4] <- 0

po[8,t,5] <- 0

po[8,t,6] <- 0

po[8,t,7] <- 0

po[9,t,1] <- 1

po[9,t,2] <- 0

po[9,t,3] <- 0

po[9,t,4] <- 0

po[9,t,5] <- 0

po[9,t,6] <- 0

po[9,t,7] <- 0

po[10,t,1] <- 1

po[10,t,2] <- 0

po[10,t,3] <- 0

po[10,t,4] <- 0

po[10,t,5] <- 0

po[10,t,6] <- 0

po[10,t,7] <- 0

\# Probabilities of states at a given occasion given states at the occasion before (see Appendix S1)

px[1,t,1] <- pSMSL[1,t]*survS[1,t]

px[1,t,2] <- pSMSL[2,t]*survS[1,t]

px[1,t,3] <- pSMSL[3,t]*survS[1,t]

px[1,t,4] <- survS[2,t]

px[1,t,5] <- survS[3,t]

px[1,t,6] <- survS[4,t]

px[1,t,7] <- survS[5,t]

px[1,t,8] <- survS[6,t]

px[1,t,9] <- survS[7,t]

px[1,t,10] <- 0

px[2,t,1] <- 0

px[2,t,2] <- (1-psiML[t])*survM[1,t]

px[2,t,3] <- psiML[t]*survM[1,t]

px[2,t,4] <- 0

px[2,t,5] <- survM[2,t]

px[2,t,6] <- survM[3,t]

px[2,t,7] <- 0

px[2,t,8] <- survM[4,t]

px[2,t,9] <- survM[5,t]

px[2,t,10] <- 0

px[3,t,1] <- 0

px[3,t,2] <- 0

px[3,t,3] <- survL[1,t]

px[3,t,4] <- 0

px[3,t,5] <- 0

px[3,t,6] <- survL[2,t]

px[3,t,7] <- 0

px[3,t,8] <- 0

px[3,t,9] <- survL[3,t]

px[3,t,10] <- 0

for (i in 4:10){

px[i,t,10] <- 1

for (j in 1:9){

px[i,t,j] <- 0

}

}

}

for (i in 1:N) # for each female

{

\# Estimated probabilities of initial states are the proportions in each state at first capture occasion

alive[i,First[i]] ~ dcat(px0[1:10])

mydata[i,First[i]] ~ dcat(po.init[alive[i,First[i]],])

for (j in (First[i]+1):Last[i])

{

\# State equations

\# Draw states at j given states at j-1

alive[i,j] ~ dcat(px[alive[i,j-1],j-1,])

\# Observation equations

\# Draw observations at j given states at j

mydata[i,j] ~ dcat(po[alive[i,j],j-1,])

}

}

#----------------------------------------------------------------

\# 2.3 Likelihood for reproductive data

#----------------------------------------------------------------

for (t in 1:(nyears-1)){

rhoS[t]<-FECUND_S[t]*R[t,1] # Small

J[t,1] ~ dpois(rhoS[t])

rhoM[t]<-FECUND_M[t]*R[t,2] # Medium

J[t,2] ~ dpois(rhoM[t])

rhoL[t]<-FECUND_L[t]*R[t,3] # Large

J[t,3] ~ dpois(rhoL[t])

}

}

",fill = TRUE)

sink()

##################################################################

\# 3 – DATA, INITIAL VALUES AND PARAMETERS MONITORED

##################################################################

\# Data

bugs.data <- list(yHS = yHS, yHM = yHM, yHL = yHL, nyears = nyears, N=n, mydata=as.matrix(mydata+1), First=e, Last=last, alphaS=rep(1,7), alphaM=rep(1,5), alphaL=rep(1,3),t ransitionS=rep(1,3), effHS=effHS, effHM=effHM, effHL=effHL, J=J[1:(nyears-1),],R=R[1:(nyears-1),])

\# Initial values

alive=mydata

for (i in 1:n) {

for (j in 1:K) {

if (j < e[i]) {alive[i,j] <- NA}

}

}

for (i in 1:n) {

if (e[i] == K) 2+2

else {

for (j in (e[i]+1):K) {

if (alive[i,j]==0 & alive[i,j-1]==1 & sum(alive[i,j:K]==2)>0) {alive[i,j] <- 1}

if (alive[i,j]==0 & alive[i,j-1]==1 & sum(alive[i,j:K]==2)==0 & sum(alive[i,j:K]==6)==0 & sum(alive[i,j:K]==3)==0) {alive[i,j] <- 1}

if (alive[i,j]==0 & alive[i,j-1]==1 & sum(alive[i,j:K]==3)>0) {alive[i,j] <- 2}

if (alive[i,j]==0 & alive[i,j-1]==1 & sum(alive[i,j:K]==3)==0 & sum(alive[i,j:K]==6)==0) {alive[i,j:K] <- 1}

if (alive[i,j]==0 & alive[i,j-1]==1 & sum(alive[i,j:K]==1)>0) {alive[i,j] <- 1}

if (alive[i,j]==0 & alive[i,j-1]==2 & sum(alive[i,j:K]==2)>0) {alive[i,j] <- 2}

if (alive[i,j]==0 & alive[i,j-1]==2 & sum(alive[i,j:K]==2)==0 & sum(alive[i,j:K]==6)==0) {alive[i,j] <- 2}

if (alive[i,j]==0 & alive[i,j-1]==2 & sum(alive[i,j:K]==3)>0) {alive[i,j] <- 2}

if (alive[i,j]==0 & sum(alive[i,j-1]==3)>0) {alive[i,j] <- 3}

if (alive[i,j]==0 & alive[i,j-1]==1 & sum(alive[i,j:K]==6)>0) {alive[i,j] <- 2}

if (alive[i,j]==0 & alive[i,j-1]==2 & sum(alive[i,j:K]==6)>0) {alive[i,j] <- 2}

}

}

}

for (i in 1:n) {

for (j in 1:K) {

if (mydata[i,j]==4 & j<K) {alive[i,(j+1):K] <- NA}

if (mydata[i,j]==5 & j<K) {alive[i,(j+1):K] <- NA}

if (mydata[i,j]==6 & j<K) {alive[i,(j+1):K] <- NA}

}

}

alive <- as.matrix(alive)

init1 <- list(alive=alive)

init2 <- list(alive=alive)

init3 <- list(alive=alive)

inits <- list(init1,init2,init3)

\# Parameters monitored

parameters <- c("sigyS", "sigyM", "sigyL","pS","pM","pL","h", "pSMSL", "psiML", "piVS", "ppS", "ppM", "ppL", "llS", "llM", "llL", “NS", "NM", "NL", "fecL", "fecM", "survS", "survM", "survL", "fec_r", "FECUND_S", "FECUND_M","FECUND_L", "BPS", "BPM", "BPL", "piOS", "SPN")

##################################################################

\# 4 – RUN THE MODEL

##################################################################

\# MCMC settings

ni <- 50000

nt <- 100

nb <- 180000

nc <- 3

\# Call JAGS from R

library(rjags)

start<-as.POSIXlt(Sys.time())

jmodel <- jags.model("ipm-prod.bug", bugs.data, inits, n.chains = nc, n.adapt = nb)

jsample <- coda.samples(jmodel, parameters, n.iter=ni, thin = nt)

end <-as.POSIXlt(Sys.time())

duration = end-start
